# Mechanics of exploration in *Drosophila melanogaster*

**DOI:** 10.1101/354795

**Authors:** Jane Loveless, Konstantinos Lagogiannis, Barbara Webb

**Affiliations:** Institute for Perception, Action, and Behaviour, School of Informatics, University of Edinburgh, 10 Crichton Street, Edinburgh EH8 9AB, Scotland, UK; MRC Centre for Developmental Neurobiology, New Hunt’s House, King’s College London, Guy’s Hospital Campus, London SE1 1UL, UK

## Abstract

The *Drosophila* larva executes a stereotypical exploratory routine that appears to consist of stochastic alternation between straight peristaltic crawling and reorientation events through lateral bending. We present a model of larval mechanics for axial and transverse motion over a planar substrate, and use it to develop a simple, reflexive neuromuscular model from physical principles. In the absence of damping and driving, the mechanics of the body produces axial travelling waves, lateral oscillations, and unpredictable, chaotic deformations. The neuromuscular system counteracts friction to recover these motion patterns, giving rise to forward and backward peristalsis in addition to turning. The model produces spontaneous exploration, even though the model nervous system has no intrinsic pattern generating or decision making ability, and neither senses nor drives bending motions. Ultimately, our model suggests a novel view of larval exploration as a deterministic superdiffusion process which is mechanistically grounded in the chaotic mechanics of the body.

## Introduction

Exploratory search is a fundamental biological behaviour, observed in most phyla. It has consequently become a focus of investigation in a number of model species, such as larval *Drosophila*, in which neurogenetic methods can provide novel insights into the underlying mechanisms. However, appropriate consideration of biomechanics is needed to understand the control problem that the animal’s nervous system needs to solve.

When placed on a planar substrate (typically, an agar-coated petri dish), the *Drosophila* larva executes a stereotypical exploratory routine [1] which appears to consist of a series of straight runs punctuated by reorientation events [2]. Straight runs are produced by laterally symmetric peristaltic compression waves, which propagate along the larval body in the same direction as overall motion (i.e. posterior-anterior waves carry the larva forwards relative to the substrate, anterior-posterior waves carry the larva backwards) [3]. Reorientation is brought about by laterally asymmetric compression and expansion of the most anterior body segments of the larva, which causes the body axis of the larva to bend [2].

Peristaltic crawling and reorientation are commonly thought to constitute discrete behavioural states, driven by distinct motor programs [2]. In exploration, it is assumed, alternation between these states occurs stochastically, allowing the larva to search its environment through an unbiased random walk [1,4–6]. The state transitions or direction and magnitude of turns can be biased by sensory input to produce taxis behaviours [4,5,7–13]. The neural circuits involved in producing the larval exploratory routine potentially lie within the ventral nerve cord (VNC), since silencing the synaptic communication within the brain and subesophageal ganglia (SOG) does not prevent substrate exploration [1]. Electrophysiological and optogenetic observations of fictive locomotion patterns within the isolated VNC [14,15] support the prevailing hypothesis that the exploratory routine is primarily a result of a centrally generated motor pattern. As such, much recent work has focused on identifying and characterising the cells and circuits within the larval VNC [16–32]. However, behaviour rarely arises entirely from central mechanisms; sensory feedback and biomechanics often play a key role [33,34] including the potential introduction of stochasticity. Indeed, thermogenetic silencing of somatosensory feedback in the larva leads to severely retarded peristalsis [35] or complete paralysis [36,37].

In line with the ethological distinctions drawn between runs and turns, computational modelling of the mechanisms underlying larval behaviour has so far focused on either peristaltic crawling or turning. An initial model based on neural populations described a possible circuit architecture and dynamics underlying the fictive peristaltic waves observed in the isolated ventral nerve cord [38]. A subsequent model described the production of peristaltic waves through interaction of sensory feedback with biomechanics, in the absence of any centrally generated motor output [39]. This model produced only forward locomotion as it incorporated strongly asymmetric substrate interaction. Recently, a model combining biomechanics, sensory feedback, and central pattern generation reproduced many features of real larval peristalsis [40]. However, this model only aimed to explain forward locomotion, and accordingly contained explicit symmetry-breaking elements in the form of posterior-anterior excitatory couplings between adjacent segments of the VNC, and posterior-anterior projections from proprioceptive sensory neurons in one segment into the next segment of the VNC. No biomechanical models of turning in the larva have yet been published, but the sensory control of reorientation behaviour has been explored in more abstract models [4,5,8,11–13,41]. No current model accounts for both peristalsis and reorientation behaviours, and no current model of peristalsis can account for both forward and backward locomotion without appealing to additional neural mechanisms.

Here we present a model of unbiased substrate exploration in the *Drosophila* larva that captures forward and backward peristalsis as well as reorientation behaviours. We provide a deterministic mathematical description of body mechanics coupled to a simple, reflexive nervous system. In contrast to previous models, our nervous system has no intrinsic pattern-generating ability [38,40,41], and does not explicitly encode discrete behavioural states or include any stochasticity [4,5,8,11–13]. Nevertheless, the model is capable of producing apparently random “sequences” of crawling and reorientation behaviours, and is able to effectively explore in a two-dimensional space. We argue that the core of this behaviour lies in the chaotic mechanical dynamics of the body, which result from an energetic coupling of axial (“peristaltic”) and transverse (“turning”) motions.

In what follows, we first outline the key components and assumptions of our model of body mechanics. We use simple physical arguments to guide the construction of a neuromuscular model capable of producing power flow into the body, and motion of the body’s centre of mass relative to the substrate. Crucially, the neuromuscular model neither senses nor drives transverse motions. In analysing the behaviour of our model, we begin by focusing on the small-amplitude, energy-conservative behaviour of the body in the absence of frictive and driving forces. In this case, the motion of the body is quasiperiodic and decomposes into a set of energetically isolated axial travelling waves and transverse standing waves. Reintroducing friction and driving forces, we demonstrate the emergence of a pair of limit cycles corresponding to forward and backward peristaltic locomotion, with no differentiation of the neural activity for the two states. We then shift focus to the behaviour of the model at large amplitudes. In this case the axial and transverse motions of the body are energetically coupled, and the conservative motion becomes chaotic. The energetic coupling allows our neuromuscular model to indirectly drive transverse motion, producing chaotic body deformations capable of driving substrate exploration. Analysis of our model supports a view of larval exploration as an (anomalous) diffusion process grounded in the deterministic chaotic mechanics of the body.

## Model specification and core assumptions

### Mechanics

To explore larval crawling and turning behaviours, we choose to describe the motion of the larval body axis (midline) in a plane parallel to the substrate (Fig 1, S1 Fig). The larval body is capable of more diverse motions including lifting/rearing [21], rolling [42], digging [43], self-righting / balancing, and denticle folding which we have recently observed to occur during peristalsis (S1 Video). However, while exploring flat surfaces, the larva displays fairly little out-of-plane motion (neither translation perpendicular to the substrate nor torsion around the body axis) and only small radial deformations [44]. Furthermore, the majority of ethological characterisations of larval exploration treat the animal as if it were executing purely planar motion [4,6,8–13,45]. A planar model is thus a reasonable abstraction for the issues addressed in this paper, i.e., the generation of peristalsis and bending.

**Fig 1.**
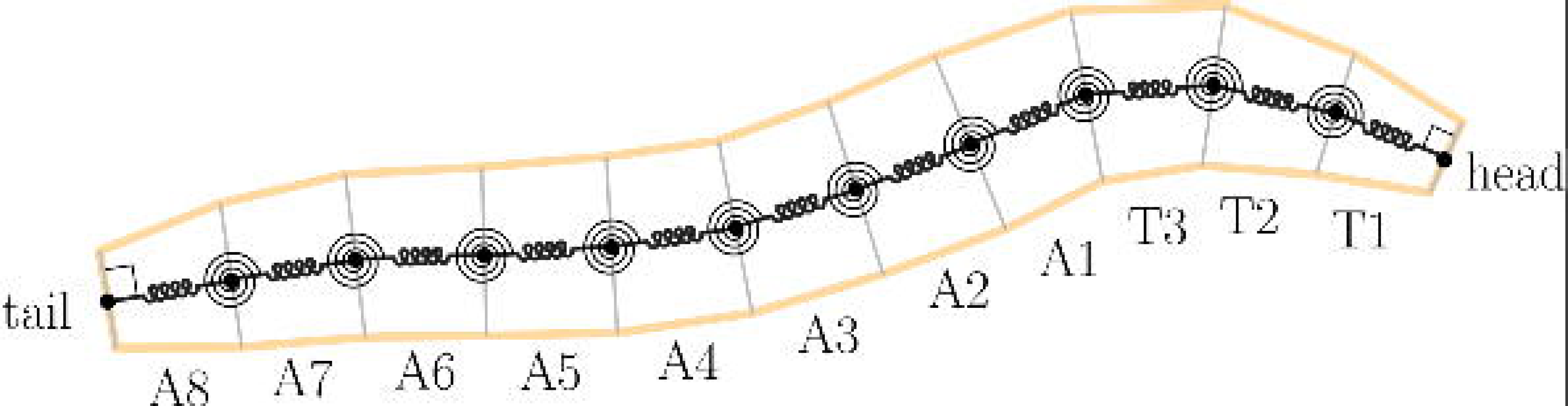
Our model of axial and transverse motion over a planar substrate. The midline of the larva is modelled as a set of discrete point masses interacting with each other via linear, damped translational and torsional springs, and with the environment via Coulomb sliding friction. We model the larva’s incompressible coelomic fluid by constraining the total length of the midline to remain constant (see main text). Quantities used to describe deformations of the body, and interaction with the substrate, are shown in S1 Fig.

The segmented anatomy of the *Drosophila* larva allows us to focus our description of the midline to a set of *N* = 12 points in the cuticle, located at the boundaries between body segments and at the head and tail extremities. We assign each point an identical mass, and measure its position and velocity relative to a two dimensional cartesian coordinate frame fixed in the substrate (the laboratory or lab frame). We therefore have *N*_*dof*_ = 2*N* = 24 mechanical degrees of freedom. We note that our assumption of a uniform mass distribution along the midline is somewhat inaccurate, since thoracic segments are smaller than abdominal segments. However, simulations with non-uniform mass distribution (not shown) give results which are qualitatively close to those presented here.

We assume that the larval body stores elastic energy in both axial compression/expansion and transverse bending, due to the presence of elastic proteins in the soft cuticle. We assume that energy is lost during motion due to viscous friction within the larva’s tissues and sliding friction between the body and the substrate. Sliding friction also allows shape changes (deformations) of the body to cause motion of the larva as a whole relative to the substrate (centre of mass motion).

Since the mechanical response of the larva’s tissues is yet to be experimentally determined, we assume a linear viscoelastic model. This is equivalent to placing linear (Hookean) translational and torsional springs in parallel with linear (Newtonian) dampers between the masses in the model, as shown in Fig 1, or to taking quadratic approximations to the elastic potential energy and viscous power loss (as in S1 Appendix). We note that the accuracy of the approximation may decrease for large deformations, in which nonlinear viscoelastic effects may become important.

As with larval tissue mechanics, there has been little experimental investigation of the forces acting between the larva and its environment. We therefore assume a simple anisotropic Coulomb sliding friction model, in which the magnitude of friction is independent of the speed of motion, but may in principle depend upon the direction of travel. This anisotropy could be thought of as representing the biased alignment of the larva’s denticle bands, or directional differences in vertical lifting or denticle folding motions which are not captured by our planar model. A mathematical formulation of our sliding friction model is given in S1 Appendix.

In addition to power losses due to friction, we also allow power flow due to muscle activation. For the sake of simplicity, we choose to allow only laterally symmetric muscle tensions. In this case, the musculature cannot directly cause bending of the midline, and can only explicitly drive axial motions. We will see later that even indirect driving of bending motion can lead to surprisingly complex behaviour.

Finally, we model the internal coelomic fluid of the larva. Given the extremely small speed of the fluid motion compared to any reasonable approximation to the speed of sound in larval coelomic fluid, we can safely approximate the fluid flow as incompressible [46]. This would ordinarily require that the volume contained within the larval cuticle remain constant. However, since we are modelling only the motion of the midline and neglecting radial deformations, we constrain the total length of the larva to remain constant. We note that this constraint is not entirely accurate to the larva, as the total length of the animal has been observed to vary during locomotion [44]. Nevertheless, for the sake of simplicity we will continue with this constraint in place, noting that this approximation has been used with success in previous work focused on peristalsis [39,40], and that there is experimental support for kinematic coupling via the internal fluid of the larva [3]. We note that we satisfy the incompressibility condition only approximately in some sections *(Model behaviour - Conservative chaos, Dissipative chaotic deformations*, and *Deterministic exploration*), by introducing an additional potential energy associated with the constraint, which produces an energetic barrier preventing large changes in the total length of the body (see S1 Appendix for details of this approximation along with specifics of the mathematical formulation of our mechanical model).

### Neuromuscular System

Let us now consider how we should use muscle activity to produce locomotion. There are two basic requirements. First, we must have power flow into the body from the musculature, so that the effects of friction may be overcome and the larva will not tend towards its equilibrium configuration. Second, we must be able to produce a net force on the centre of mass of the larva, so that it can accelerate as a whole relative to the lab frame. Note that in this section, we motivate the neural circuits in the model from this purely functional point of view, but will present relevant biological evidence in the discussion.

To satisfy the first criterion, let us examine the flow of power into the body due to the action of the musculature

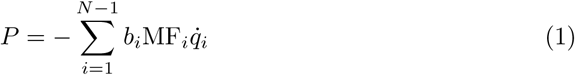

Here, *q*_*i*_ describes the change in length of the *i*’th body segment away from its equilibrium length, 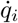 is the rate of expansion of the *i*’th body segment, *b*_*i*_ is a (positive) gain parameter, MF_*i*_ is a (positive) dimensionless control variable representing muscle activation, and the product *b*_*i*_MF_*i*_ is the total axial tension across the *i*’th body segment. From this expression, it is clear that if we produce muscle tensions (MF_*i*_> 0) only while segments are shortening 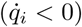, we will always have positive power flow into the body (*P* > 0). This is a mathematical statement of the requirement for the larva’s muscles to function as *motors* during locomotion, rather than as springs, brakes, or struts [33].

A simple way to fulfil this condition is to introduce a segmentally localised reflex circuit (Fig 2, [39]). We place a single sensory neuron in each segment which activates when that segment is compressing 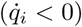. Each sensory neuron then projects an excitatory connection onto a local motor neuron, which in turn projects to a muscle fibre within the same segment. Assuming for now that there are no other influences on the motor neurons, so that sensory activation implies local motor neuron activation, segmental shortening will produce an immediate muscle tension serving to amplify compression of the segment and thus counteract frictive energy losses.

**Fig 2.**
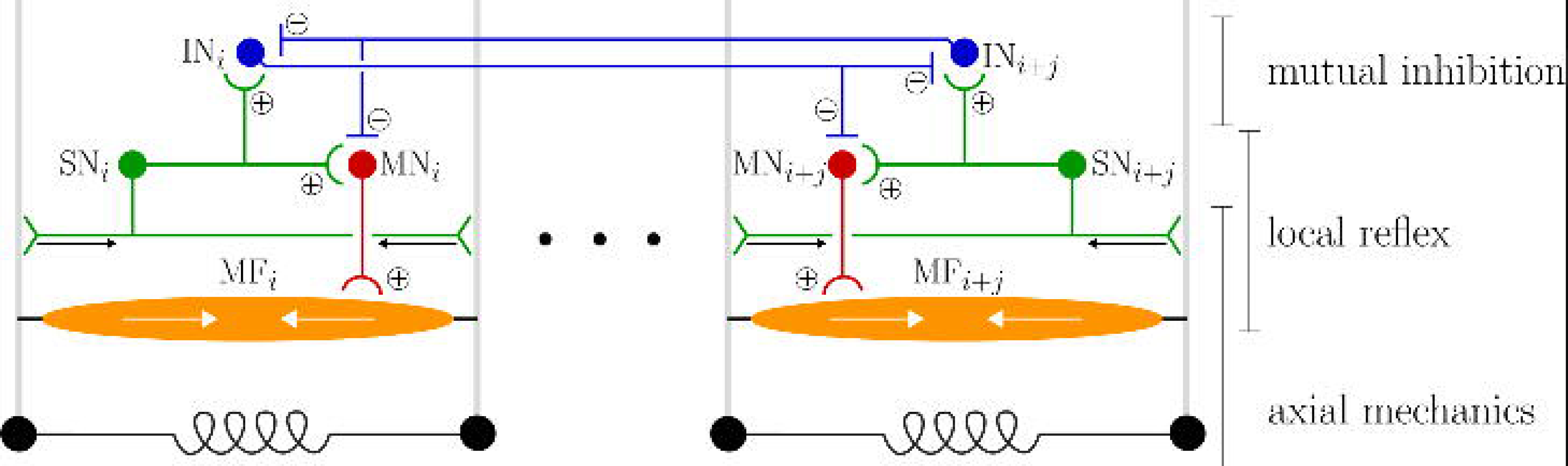
The neuromuscular model. A local reflex amplifies motion via positive feedback: sensory neurons SN activate during segmental shortening, exciting motor neurons MN, and causing muscle fibre activation MF which accelerates shortening. Reflexes in distant segments *i* and *i* + *j* (|*j*| > 1) mutually inhibit one another via interneurons IN. This limits the number of moving segments to allow centre of mass motion (see text).

Let us now consider the second criterion for peristaltic locomotion. Assuming all segment boundaries are of equal mass, the force on the centre of mass of the larva is proportional to the sum of the forces acting on each segment boundary, i.e.

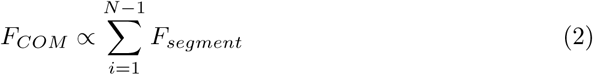

Newton’s third law tells us that any forces of interaction between segment boundaries (i.e. viscoelastic and muscle forces) must be of equal magnitude and opposite direction, so that they cancel in this summation and we are left only with contributions arising from substrate interaction. If the motion of the body is such that some number *n*_*f*_ of segments move forward at a given time, against a frictional force -*μ*_*f*_, while *n*_*b*_ segments remain anchored or move backward, experiencing a frictional force *μ*_*b*_, then the summation becomes

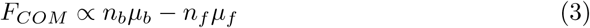

In the limiting case of isotropic (direction-independent) substrate interaction we have *μ*_*b*_ = *μ*_*f*_, and this expression tells us that the centre of mass will accelerate in the forward direction only when there are less segments moving forward than are moving backward or anchored to the substrate. Similarly, moving a small number of segments backward while the others remain anchored will result in backward acceleration of the centre of mass. Therefore, if the animal is to move relative to its substrate, it must ensure that only a limited number of its segments move in the overall direction of travel at a given time (indeed, this matches observations of the real larva [3,22]). A more lengthy exposition of this requirement on limbless crawling behaviours can be found in [47].

We fulfil the requirement for a small number of moving segments by introducing mutually inhibitory interactions between the segmentally localised reflex circuits (Fig 2). We add a single inhibitory interneuron within each segment. When the sensory neuron within the local reflex activates, it excites this interneuron, which then strongly inhibits the motor neurons and inhibitory interneurons in non-adjacent segments, effectively turning off the local reflexes in distant neighbours. Adjacent segments do not inhibit each other in our model, allowing reflex activity to track mechanical disturbances as they propagate from one segment to the next. We comment on the plausibility of this feature of our model, given the experimental observation of nearest-neighbour inhibitory connections in the larval ventral nerve cord [28], in the discussion. Similarly, the head and tail segments do not inhibit each other, which permits peristaltic waves to be (mechanically) reinitiated at one extremity as they terminate at the other. This effectively introduces a ring-like topology into the neural model, matching our model of axial mechanics which couples head and tail motion through the total length constraint [39].

We now have a neuromuscular model consisting of four cell types repeated in each segment - sensory neurons, inhibitory interneurons, motor neurons, and muscle fibres. For the sake of simplicity we model all neurons as having a binary activation state governed by the algebraic relation

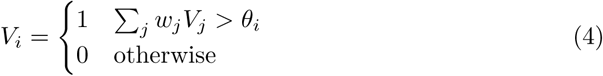

where *V*_*i*_ is the activation of the *i*’th cell, *θ*_*i*_ is its activation threshold, *V*_*j*_ is the activation of the *j’*th presynaptic cell, and *w*_*j*_ is the associated synaptic weight. Numerical values for the weights and thresholds used in our model are given in S1 Table, supplemental. Note that the muscle tension over a segment either vanishes (when the muscle fibre is in the inactive state) or has fixed magnitude *b*_*i*_ (when the muscle fibre is activated by local sensory feedback). For this reason we refer to *b*_*i*_ as the *reflex gain.*

Our choice to neglect neural dynamics is based on the large difference in timescales between the neural and mechanical dynamics. Any rapid changes in the muscle tension across a segment (due to motor neuron spiking) will be averaged out by the body’s mechanics. Note that the lack of neural dynamics in our model immediately rules out central pattern generation.

To summarise, the neural model we have constructed can be seen as consisting of two parts, a segmentally repeating local reflex and a mutual inhibition circuit acting between non-adjacent reflexes. The local reflex is constructed so that muscles will act as motors, amplifying segmental compressions and counteracting friction. The mutual inhibition circuit couples distant reflexes to allow only localised amplification. By limiting the number of moving segments, this should ensure that the model larva can produce a net force on its centre of mass.

## Model behaviour

### Larva-like axial compression waves and lateral oscillations result from conservative mechanics

One of the advantages of grounding our model of larval exploration in the body’s physics is that we are now able to apply powerful analytical results from classical mechanics in order to understand the model’s behaviour. In this section we attempt to elucidate the naturally preferred motions of the larva by focusing our attention on the conservative mechanics of the body while neglecting friction forces, which would cause all motion to stop, and driving forces, which might impose a particular pattern of motion.

In this case, the general character of motion is specified by the Liouville-Arnold integrability theorem. This theorem asks us to look for a set of conserved quantities associated with a mechanical system, which remain unchanged as the system moves (energy, momentum, and angular momentum are examples of some commonly conserved quantities). If we can find a number of these quantities equal to the number of mechanical degrees of freedom in our model, then the theorem tells us that the motion of the body is *integrable* - it can be expressed analytically, and must be either periodic or quasiperiodic. If there are not enough conserved quantities, then the system is said to be *nonintegrable.* In this case the motion is much more complicated and will be chaotic for some initial conditions. These chaotic motions do not permit analytical expression and must be approximated through simulation.

In this section, we explicitly seek a case for which there is a “full set” of conserved quantities (we provide only major results here, for detailed derivations see S2 Appendix). We begin by restricting ourselves to considering only small deformations of the larval midline, in the case where all segments are of identical axial stiffness *k*_*a*_, transverse stiffness *k*_*t*_, mass *m*, and length *l*. Under these assumptions, the total mechanical energy of the body may be written

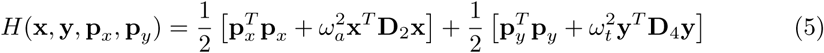

where **x** and **y** are vectors giving the displacement of each segment boundary along the body axis and perpendicular to the body axis, respectively, **p**_*x*_ and **p**_*y*_ give the translational momentum associated with each direction, **D**_2_ and **D**_4_ are difference matrices arising from a Taylor series expansion of our model’s potential energy (see S2 Appendix), and 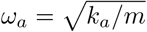 and 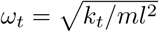 are characteristic axial and transverse frequency scales. By making a linear change of coordinates

{**X**,**y**,**P***x*,**P***y*}→{**X**,**Y**,**P_*X*_**,**P_*Y*_**}to the eigenbasis of **D**_2_ and **D**_4_ (see S2 Appendix) this simplifies to

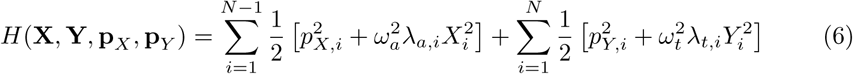

where λ_*a,i*_ and λ_*t,i*_ are eigenvalues associated with the coordinate transformation. This expression is a sum of component energies, each of which is independently conserved. The Liouville-Arnold theorem immediately tells us that the motion of the body must be (quasi)periodic in the case of conservative small deformations. Indeed, the energy associated with each of the new coordinates *X_i_,Y_i_* is in the form of a harmonic oscillator, and each coordinate executes pure sinusoidal oscillations. By transforming back to the original coordinates **x, y** we obtain a set of collective motions (modes) of the body which are energetically isolated and have a sinusoidal time dependence, corresponding to axial and transverse standing waves. We will refer to the *X_i_,Y_i_* as modal coordinates since they describe the time dependence of each of the collective motions.

Each transverse standing wave corresponds to a periodic lateral oscillation of the body, with a unique frequency given by 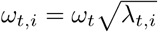. We determined these frequencies numerically, along with the spatial components of the lowest frequency standing waves (Fig 3A). These can be seen to match the eigenmaggot shapes extracted from observations of unbiased larval behaviour [48].

**Fig 3.**
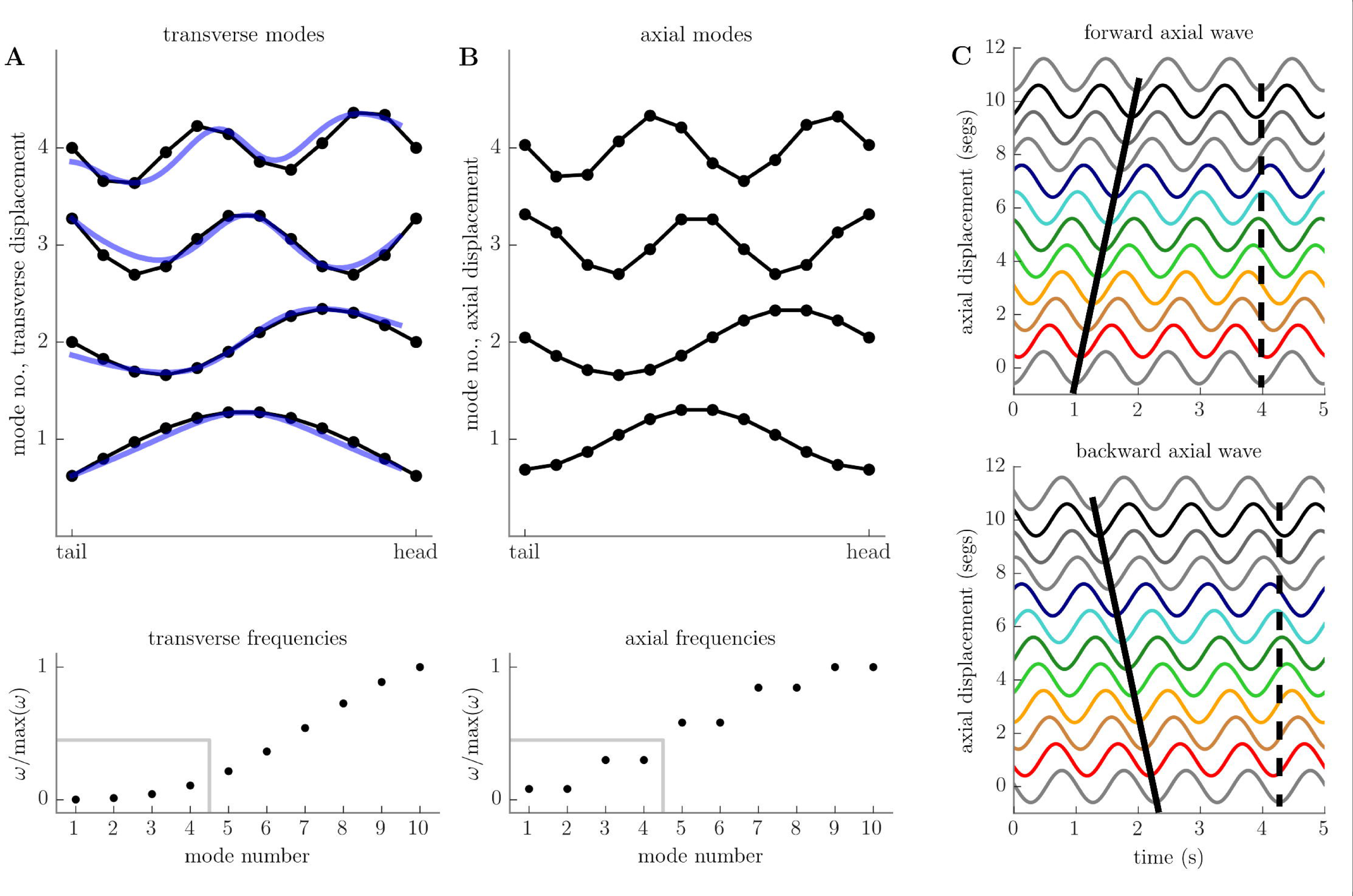
Conservative, small-amplitude motions of the body decompose into a set of axial and transverse standing waves. A: spatial component of first four transverse standing waves (top, black) compared to first four experimentally determined eigenmaggots [48] (top, blue), with natural frequencies of oscillation (bottom). B: spatial component of first four axial standing waves (top), with natural frequencies of oscillation (bottom). Note that axial standing waves come in pairs with identical frequency. C: Pairs of axial standing waves can be combined to produce forward-propagating (top, solid black line) and backward-propagating (bottom, solid black line) travelling waves. Head and tail extremities move in phase (dashed black line) due to our total length constraint (see text), reminiscent of the “visceral pistoning” observed in the real animal [3].

The axial standing waves correspond to oscillating patterns of segmental compression and expansion. Unlike the transverse waves, the axial waves come in pairs with identical frequencies of oscillation. We were able to analytically determine the frequency of oscillation of the i’th pair of standing waves to be

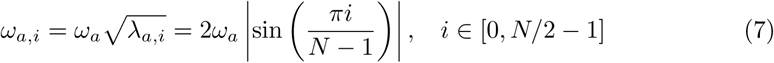

The spatial components of the axial standing waves could also be determined analytically

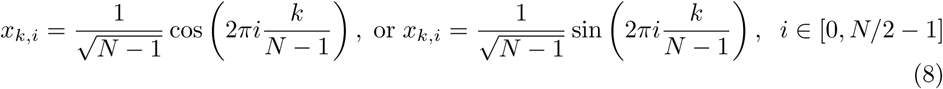

Where *x*_*k i*_ is the displacement of the *k*’th segment boundary for the *i*’th pair of standing waves. We plot the axial frequencies *ω*_*a,i*_ and spatial components *x*_*k,i*_ in Fig 3B. The degeneracy in the axial oscillation frequencies allows us to combine the axial standing waves with a ±90° relative phase shift to form pairs of forward and backward travelling wave solutions, given by

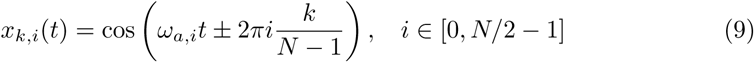

where *x*_*k,i*_ (t) gives the displacement of the *k*’th segment boundary as a function of time for the *i*’th pair of travelling waves. The choice of a plus or minus sign corresponds to the choice between forward or backward wave propagation. We plot the lowest frequency pair of axial travelling wave solutions in Fig 3C, and directly visualise the synthesis of travelling wave solutions from standing wave solutions in S2 Video.

To summarise, in this section we have shown that for the case of conservative, small oscillations, the motion of the body may be decomposed into a combination of transverse standing waves and axial travelling waves. This is of clear relevance to understanding the behaviour of the larva, which moves across its substrate by means of axial peristaltic waves while reorienting using lateral oscillations. Indeed, the transverse modes of oscillation that we have derived here match principal components of bending computed from actual larval behaviour [48]. We will now focus on the small-amplitude motion of the body in the presence of energy dissipation due to friction and driving forces.

### Forward and backward peristaltic locomotion can be obtained from simple reflexes

Reintroducing friction will clearly lead the motions described above to eventually terminate due to energy dissipation, unless opposed by transfer of power. In a previous section *(Model specification and core assumptions - Neuromuscular system*), we introduced a neuromuscular system to produce power flow into the body, but as described, it can only directly transfer power into the axial degrees of freedom. In the small deformation model we have just analysed, the axial and transverse degrees of freedom are energetically decoupled. It follows that transverse friction is unopposed and any transverse motion must eventually terminate in the case of small deformations. In this section we will therefore focus only on the axial degrees of freedom, which correspond to the peristaltic locomotion of the larva.

In Fig 4, we show the effect of coupling our neuromuscular model to the axial mechanics. For small reflex gains, the power flow into the body from the musculature is too low to effectively counteract frictive losses and the larva tends towards its passive equilibrium state, in which there is complete absence of motion. However, increasing reflex gain past a certain critical value leads to the emergence of long-term behaviours in which the larva remains in motion, away from its passive equilibrium. These motions correspond to forward and backward locomotion, driven by forward and backward propagating compression waves (see below), as predicted from our earlier description of the conservative motions of the body, and as observed in the real larva [3]. The qualitative changes in behaviour that occur as reflex gain is varied are depicted in Fig 4A, where we have measured the long-term centre of mass momentum of the larva, along with the long-term relative phase of the lowest frequency modal coordinates.

**Fig 4.**
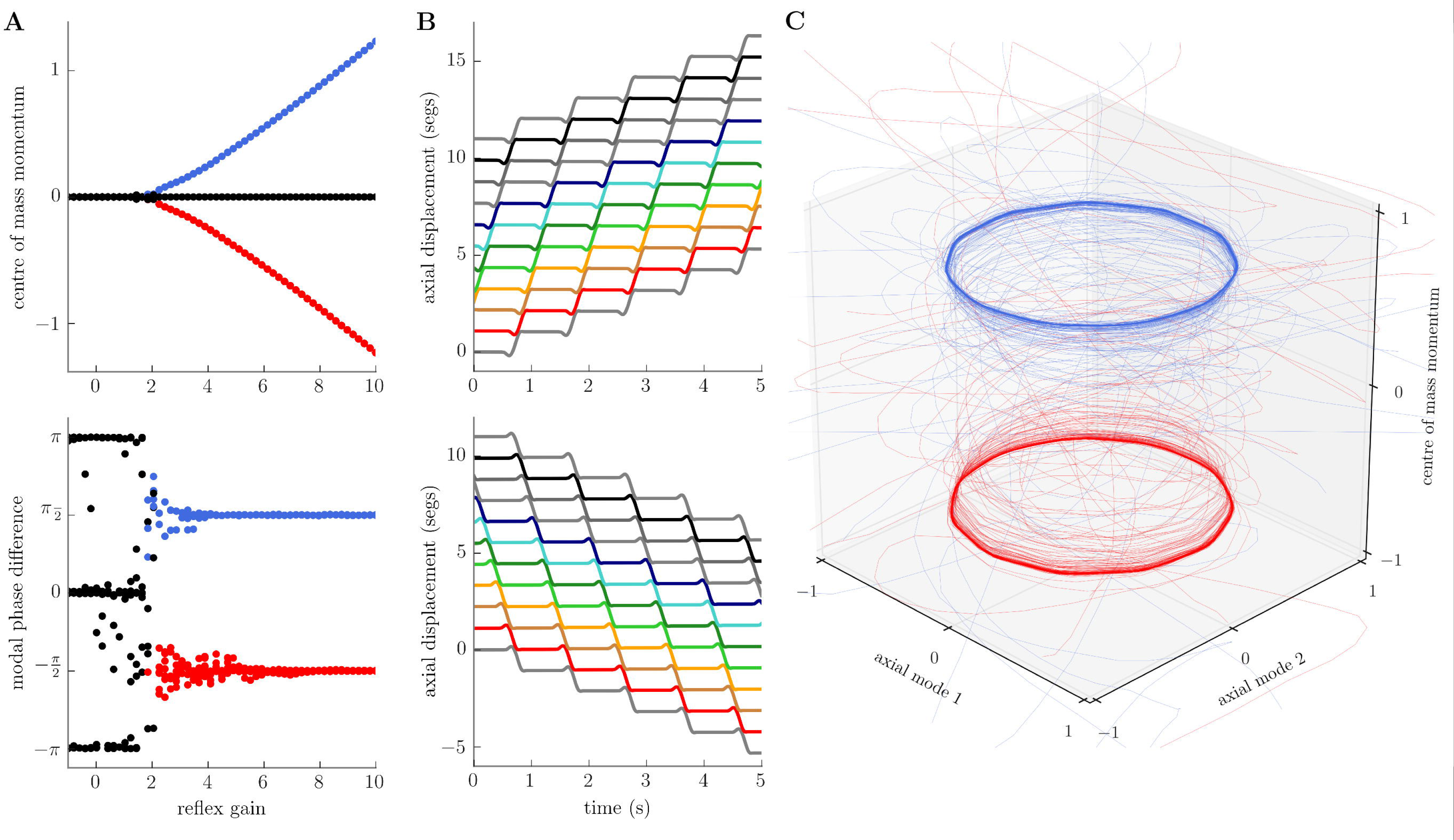
Emergence of limit cycles for forward and backward locomotion in the dissipative, small-amplitude model. A: as reflex gain is increased past a critical point, the model larva attains a positive or negative long-term average centre of mass momentum (top, red and blue lines), signifying continuous forward or backward motion relative to the substrate, and a *±*π*/2* relative phase difference between the two lowest frequency axial standing wave modes (bottom, red and blue lines), signifying the presence of forward- or backward-propagating axial travelling waves. B: trajectories of individual point masses in the model for forward (top) or backward (bottom) locomotion (see S2 Fig for corresponding neural state). C: projection of model trajectories onto the lowest frequency axial modes and the centre of mass momentum reveals a pair of (putative) stable limit cycles for forward (blue) and backward (red) locomotion. Parameters used to generate this figure are given in S2 Table.

For low reflex gains the centre of mass momentum tends to 0 as the body comes to a stop and enters a passive equilibrium state. The relative phase of the low frequency modal coordinates tends to either 0 or 180 degrees, which also corresponds to a loss of momentum. For larger values of reflex gain, the total momentum is either positive, zero, or negative. Positive and negative total momentum represent forward and backward locomotion, respectively, while zero momentum corresponds to two unstable cases which we discuss below. The relative phase of the lowest frequency modal coordinates tends to ±90° at high reflex gains, corresponding to the presence of forward- or backward-propagating compression waves (see previous section). As in the larva [1,3], forward-propagating waves drive forward locomotion while backward-propagating waves drive backward locomotion (Fig 4B).

We believe that these behaviours arise from the production of a pair of limit cycle attractors in the system’s phase space, which we visualise in figure Fig 4C by projecting the system state onto the lowest frequency pair of axial modes, and plotting the associated modal coordinates along with the centre of mass momentum. Since wave motion implies that pairs of modal coordinates should perform pure sinusoidal oscillations with equal amplitude and frequency, and a ±90^°^ relative phase shift (see previous section and S2 Appendix), the travelling wave trajectories of the system become circles in this coordinate system (see discussion of Lissajous figures, [49]). Forward and backward locomotion can then be distinguished by the momentum of the centre of mass.

In S2 Fig we show the neural state of the model larva during forwards and backwards locomotion. As expected given our previous exposition, we observe waves of activity in the nervous system which track the mechanical waves propagating through the body. Our sensory neurons also show a second, brief period of activation following propagation of the mechanical wave caused by a slight compression that occurs as segments return to equilibrium. This activity is “cancelled out” by the mutual inhibition circuit, so that motor neurons do not exhibit a secondary burst of activity.

The larva has zero long-term total momentum in the presence of large reflex gain in only two cases, both of which are highly unstable. First, if we start the larva so that it is already in its passive equilibrium state, so that no relative motion of segment boundaries occurs, it is obvious that there will be no activation of local reflexes and the larva will not spontaneously move out of equilibrium. The second case corresponds to a pure axial standing wave. In this case the larva is divided into two regions by nodal points where the axial displacement is zero. During the first half-cycle of the standing wave, one region will experience compression while the other experiences expansion, and in the second half-cycle these roles will reverse. The neural circuit we have constructed can amplify compression during both half-cycles since they are separated by a configuration in which no compression or expansion occurs, and this allows the entire nervous system to become inactive and “reset”. Since these behaviours are extremely unstable and require very specific initial conditions to be observed, we have not visualised them here.

While the mutually inhibitory connections in our model are not required for the propagation of locomotor waves, which can be maintained entirely by local reflexes [39], these connections do greatly enhance stability. In the absence of the mutual inhibition circuit, small mechanical disturbances in any stationary body segments can be amplified, giving rise to multiple compression waves which travel through the body simultaneously (not shown). If this instability produces an equal number of forward and backward moving segments then overall motion of the larva relative to the substrate will stop, in line with the argument presented earlier. We have also observed that roughly symmetrical substrate interaction is required to produce both forward and backward locomotion in our model. If friction is too strongly anisotropic, then locomotion can only occur in one direction regardless of the direction of wave propagation (not shown).

It is worth noting that the axial model presented in this section does display discrete behavioural states. However, there are no explicit decisions regarding which behavioural states to enter, since the particular neural states occupied during forwards and backwards locomotion are indistinguishable. Rather, both the apparent decision and the eventual direction of travel can only be understood by examining the dynamics and mechanical state of the body.

### Conservative chaos from mechanical coupling

Having successfully produced peristaltic locomotion using our model, we will now turn our attention to the larva’s turning behaviours. As before, we will start from physical principles. In a previous section (*Model behaviour - Conservative axial compression waves and transverse oscillations*) we showed that, for the case of conservative small oscillations, transverse motions of the body were energetically decoupled from axial motions, and could be decomposed into a set of periodic standing waves. We will first extend our previous analysis to the case of energy-conservative, large amplitude motions in the absence of damping and driving; and then in the following section consider the impact of energy dissipation and the neuromuscular system on transverse motion,

To keep our presentation simple and allow visualisation of model trajectories, we will focus on a reduced number of the mechanical degrees of freedom. Namely, we will examine the bending angle *φ* and axial stretch *q* of the head segment (Fig 5A). We introduce an amplitude parameter ϵ by making the substitutions *q*→ ϵ*q* and *φ* → ϵ*φ*, so that the total mechanical energy of the head may be written in nondimensional form as (see S4 Appendix)

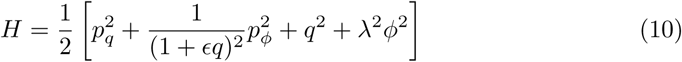

where *p*_*q*_, *pφ* are the radial and angular momentum of the head mass, and we have scaled the time axis of the model so that the natural frequency of axial oscillation is unity. The parameter 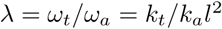 is the ratio of transverse and axial frequencies.

**Fig 5.**
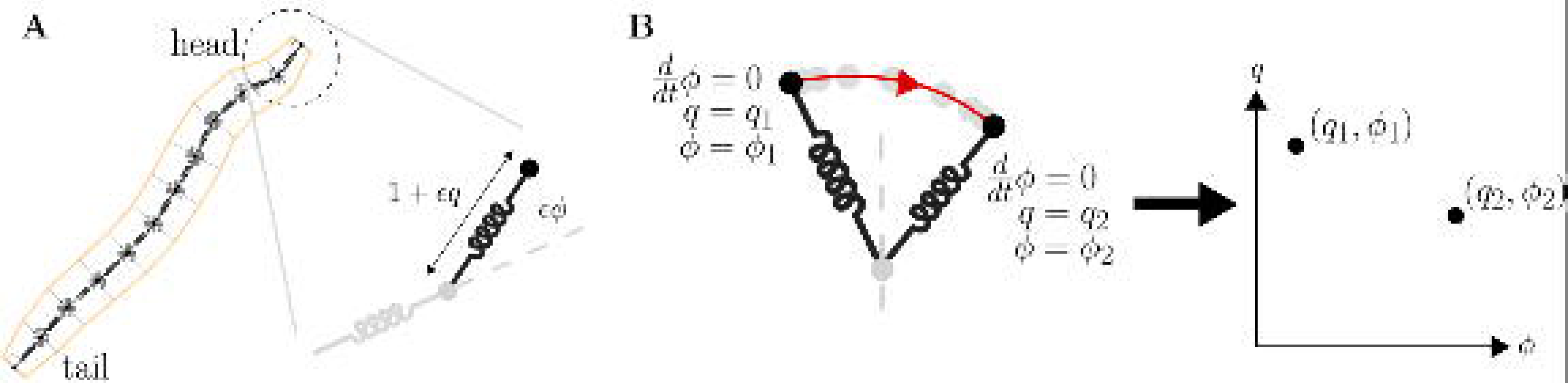
A reduced model of large amplitude motion. A: we focus on the conservative dynamics of the head’s strain *q* and bend *φ* coordinates as amplitude ϵ is varied. B: head trajectories are visualised by Poincare section, in which the head’s configuration *q, φ* is plotted at successive turning points of the transverse bending motion (at which angular velocity vanishes, d*φ* /dt = 0).

In the case of small oscillations, i.e. ϵ → 0, the mechanical energy reduces to the simpler expression

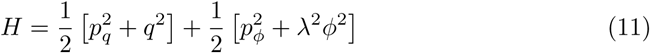

which is clearly a sum of independent axial and transverse energies. These energies are individually conserved, so that the Liouville-Arnold theorem applies, and the motion of the head is integrable and (quasi)periodic. This is in clear agreement with our earlier results. For the more general case of large amplitude motion (ϵ > 0), we may have in principle only a single conserved quantity - the total energy of the system. Indeed, it should be clear from the presence of a “mixed” term in the mechanical energy (Eq 10) that the axial and transverse motions are energetically coupled at large amplitudes, so that the individual energies associated with each motion are no longer independently conserved. Given that we have only one conserved quantity for a two degree of freedom system, we can no longer rely on the Liouville-Arnold theorem to prove (quasi)periodicity of the motion, and must accept that the system’s behaviour may be chaotic.

To investigate this possibility further, we first note that conservation of energy implies that flow within the four dimensional phase space must be constrained to lie on the energy surface given implicitly by the relation *H(q,φ,p_q_,p_φ_)* = E. Therefore, given a particular value E for the total energy, the system dynamics becomes three dimensional. This allows us to visualise the behaviour of the system by plotting the points at which trajectories intersect a two-dimensional Poincare section [50,51]. We define our Poincare section by the condition that the angular momentum vanishes *p*_*φ*_ = 0 (equivalently, angular velocity vanishes d*φ*/dt = 0), and plot successive crossings of the section as points in the *q, φ* plane. This allows us to intuitively interpret points in the Poincare section as configurations of the head at successive turning points (extrema) in the transverse motion (Fig 5B).

In what follows, we set the total energy to be 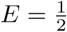 so that when ϵ = 1 we can in principle obtain complete compression of the head (*q* = −1). We choose to set 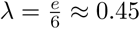, giving an irrational frequency ratio. This loosely matches observations of the real larva in which the frequency of transverse oscillations is approximately half that of axial oscillations but does not satisfy an exact (rational) resonance relationship [41]. The results we obtain with these parameters do not differ much from results for other energies or other frequency ratios, including resonant relationships (not shown).

Poincare plots for the cases ϵ → 0 and 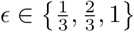 are shown in Fig 6. When ϵ → 0 (Fig 6A), conservation of transverse energy implies that the turning points of the transverse motion must remain constant. The fact that the frequency ratio λ is irrational implies that the overall motion is quasiperiodic, and the values of *q* obtained at successive transverse turning points should not repeat. In accordance with these observations, the Poincare section for ϵ → 0 consists of a series of verticle lines (Fig 6Ai). For 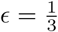 the Poincare plot becomes distorted, but the majority of trajectories still trace out one-dimensional curves in the section (Fig 6Bi), which is indicative of persistent quasiperiodic behaviour. At 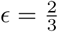 the Poincare plot appears qualitatively different. There is now a large region of what appears to be “noise”, indicating that the configuration of the head at successive transverse turning points has become unpredictable. This is a clear signature of deterministic chaos. There are, however, several regions of the section indicative of (quasi)periodic behaviour. These appear as one-dimensional curves or discrete points in the Poincare section (Fig 6Ci). At ϵ = 1, the region of the Poincare plot occupied by chaos has expanded, although there still appear to be some regions corresponding to (quasi)periodic behaviour (Fig 6Di). These results qualitatively agree with the Kolmogorov-Arnold-Moser theorem [49], which tells us that quasiperiodic behaviour should persist under small nonintegrable (chaotic) perturbations of an integrable Hamiltonian, and that the region of phase space corresponding to chaotic behaviour should grow with the perturbation size (in our case, the perturbation size corresponds to the amplitude of motion e). However, our model as presented here does not formally meet the requirements of this theorem (see S4 Appendix).

**Fig 6.**
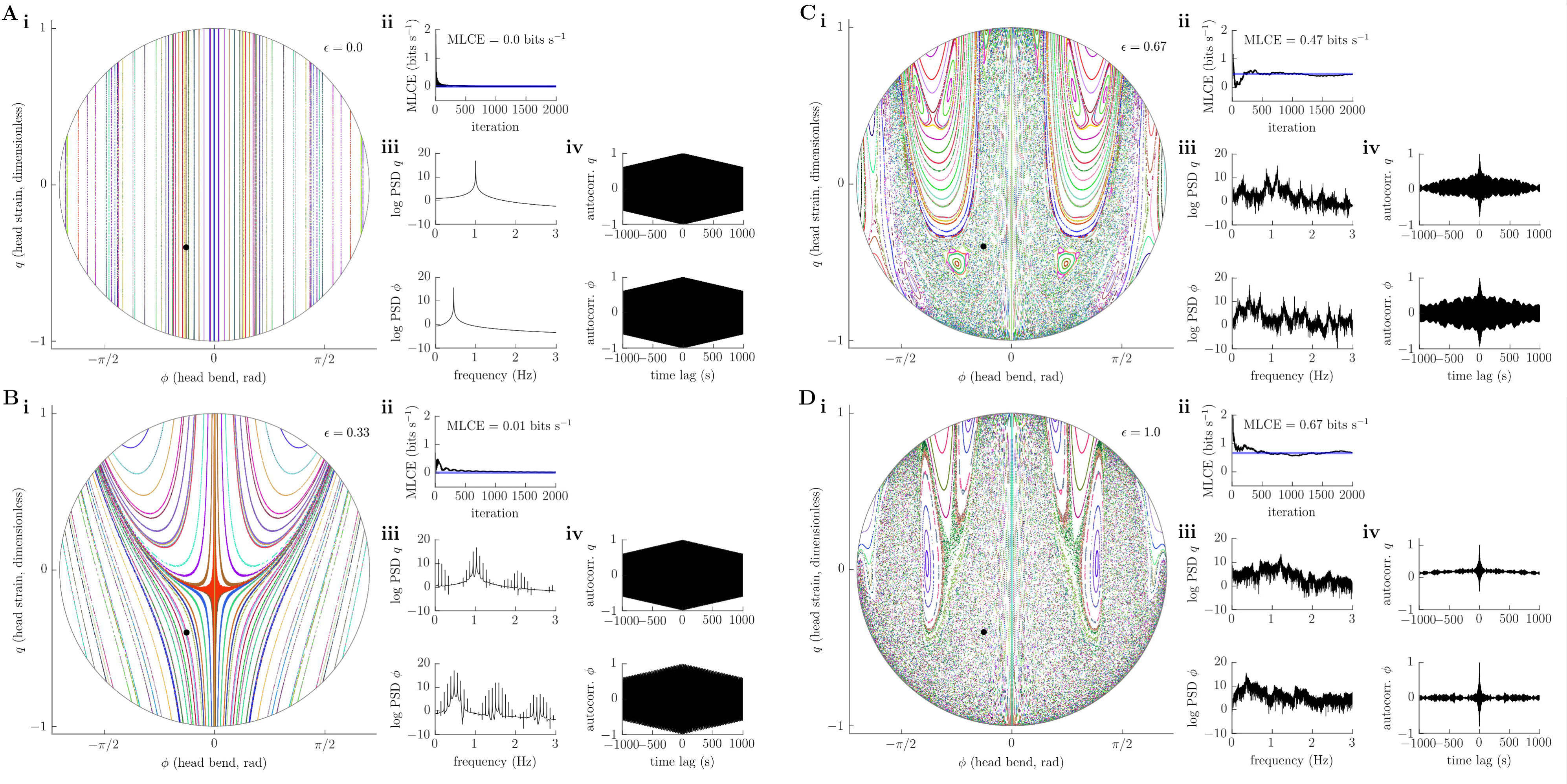
Emergence of deterministic chaos in the conservative head dynamics as amplitude of motion is increased. A, B: for small amplitudes (ϵ → 0, ϵ = 1/3), Poincare section shows quasiperiodic head oscillations (i), while the maximum Lyapunov characteristic exponent (MLCE), which quantifies the dominant rate of separation of nearby phase trajectories, converges to ~ Obits s^−1^ (ii), the power spectra of head stretch *q* and bend *φ* coordinates show clear peaks with little “noise” component (iii), and autocorrelations of these variables decay linearly (iv). These results betray non-chaotic, quasiperiodic oscillations for small amplitudes. C, D: for large amplitudes (ϵ = 2/3, ϵ = 1), the Poincare section contains a large chaotic sea (i), while the MLCE converges to a positive value (ii), power spectra become “noisy” (iii), and autocorrelations decay rapidly (iv). These results strongly suggest the existence of deterministic chaotic head dynamics for large amplitudes. MLCE, power spectra, and autocorrelations were computed for initial conditions shown by black dot in panel i. Parameters used to generate this figure are detailed in the main text, and reported in S3 Table.

Analysis by Poincare section provides an invaluable method to determine the character of overall system behaviour by direct visualisation of trajectories, but cannot be applied to higher dimensional systems. This is problematic, since we would like to assess the existence of chaos beyond our reduced model of the larva’s head. We therefore deployed a series of other methods which are possibly less reliable than the method of Poincare section but can be applied equally well to higher dimensional systems. These included estimation of the maximal Lyapunov characteristic exponent (MLCE) for the dynamics along with calculation of the power spectrum and autocorrelation of internal variables [50–52]. The MLCE can be thought of as quantifying the rate of separation of nearby trajectories, or, equivalently, the rate at which information is generated by the system being analysed [53]. A positive MLCE is generally considered a good indicator of chaotic behaviour. The power spectrum of a periodic or quasiperiodic process should consist of a “clean” set of discriminable peaks, whereas that of a chaotic process should appear “noisy” and contain power across a wide range of frequencies. Meanwhile, the autocorrelation of a periodic or quasiperiodic process should show a strong oscillatory component with an envelope that decays linearly with time, while that of a chaotic process should show a much quicker decay, similar to a coloured noise process. In Fig 6 we plot these measures at each value of e, for a trajectory starting with initial conditions indicated on the corresponding Poincare plot by a filled black circle. These measures confirm increasingly chaotic behaviour as the amplitude e increases, in agreement with our Poincare analysis. In Fig 7 we show a solution including all degrees of freedom in our conservative mechanical model (i.e. not just those of the head). The trajectory of individual segments relative to the substrate appears qualitatively irregular, while the indirect measures we introduced above (MLCE, power spectrum, autocorrelation) are all indicative of deterministic chaotic behaviour.

**Fig 7.**
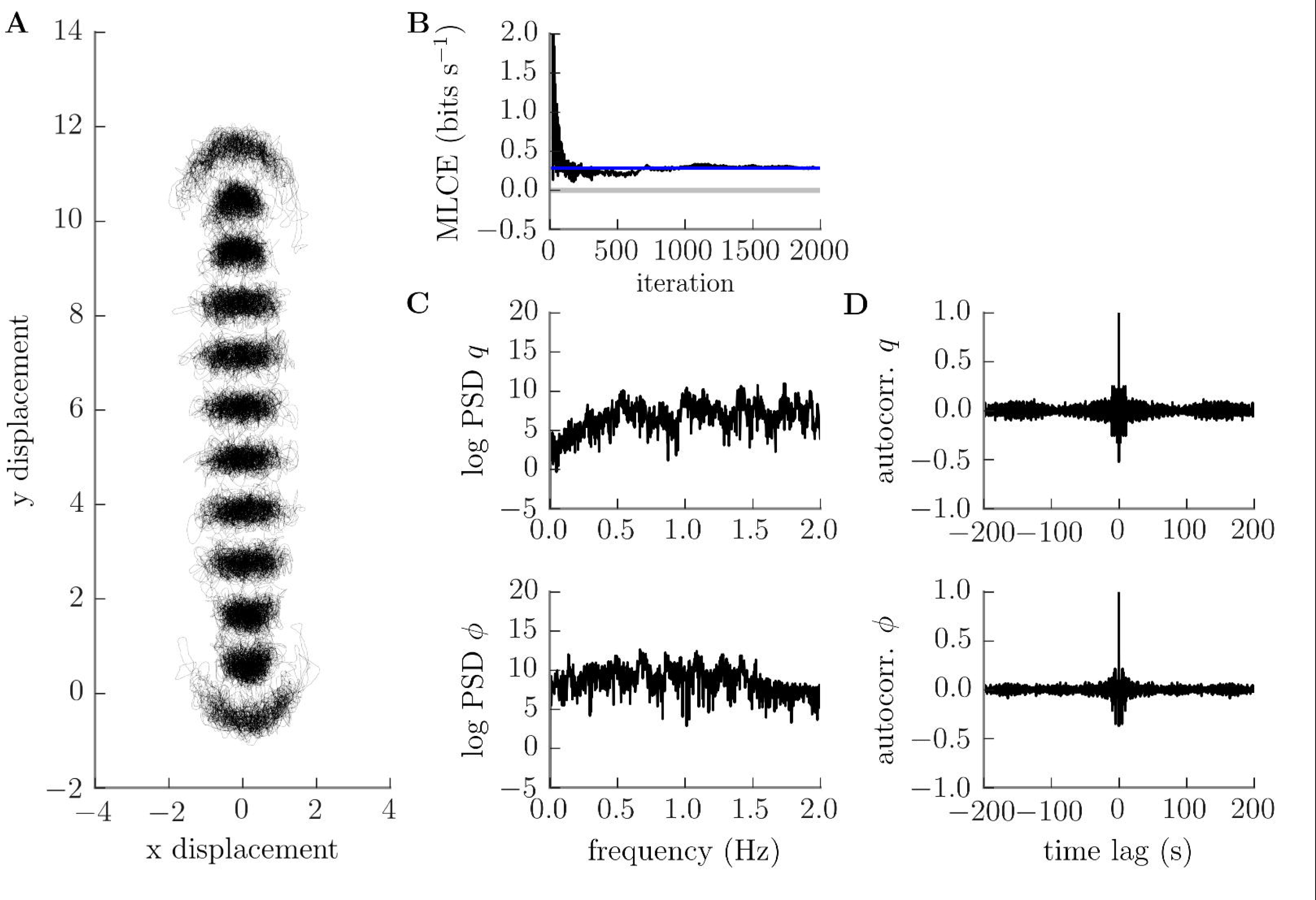
Conservative planar motion of the body is chaotic at large amplitudes. A: trajectories of individual segment boundaries appear qualitatively irregular, B: our estimate of the maximum Lyapunov characteristic exponent converges to a positive value, C: power spectra of head stretch *q* and bend *φ* show a strong “noise” component, and D: their autocorrelations decay rapidly. All are indicators of deterministic chaos. Parameters used to generate this figure are given in S4 Table.

### Spontaneous turning and reversals require no additional control

We will now reintroduce dissipative effects into our model of larval motion in the plane. We previously saw that conservative mechanics predicted axial travelling waves and transverse oscillations. These were lost in the presence of friction, but the axial travelling waves could be recovered with the addition of a neuromuscular system designed to selectively counteract frictive effects. We have now seen that conservative mechanics predicts chaotic planar motion. Although our neuromuscular model transfers power only into the axial degrees of freedom, we recall from the previous section that axial and transverse motions are energetically coupled at large amplitudes. We therefore tentatively expect that we may be able to recover the complete chaotic planar motion without requiring any additional mechanism for direct neuromuscular power transfer into tranverse motion.

To investigate whether our dissipative planar model shows chaotic behaviour, we ran *n* = 1000 simulations starting from almost identical initial conditions (euclidean distance between initial mechanical state vectors <10^−7^, with no initial neural activity) and identical parameters (see S5 Table). We can indeed observe that the simulated larva perform peristalsis with interspersed bending of the body (turns), and that the fully deterministic system produces apparently random turning such that the simulations rapidly diverge (S3 Video). Since most working definitions of chaos require strictly bounded dynamics, we here restrict our analysis to the coordinates describing deformation of the body (segmental stretches and bending angles), ignoring motions of, or overall rotations about, the centre of mass (i.e., the trajectory through space of the body, which we will analyse in the following section).

Qualitatively, the deformations of the large amplitude dissipative model appear irregular (Fig 8A). However, there are persistent features reminiscent of the ordered small-amplitude behaviour described in previous sections. In particular, there are clear axial travelling waves and lateral oscillations. However, whereas forward- and backward-propagating axial waves previously corresponded to stable limit behaviours, the large amplitude system appears to go through occasional “transitions” between these behaviours. In addition, apparently spontaneous large bends appear occasionally, suggesting that the neuromuscular system is effectively driving transverse motion.

**Fig 8.**
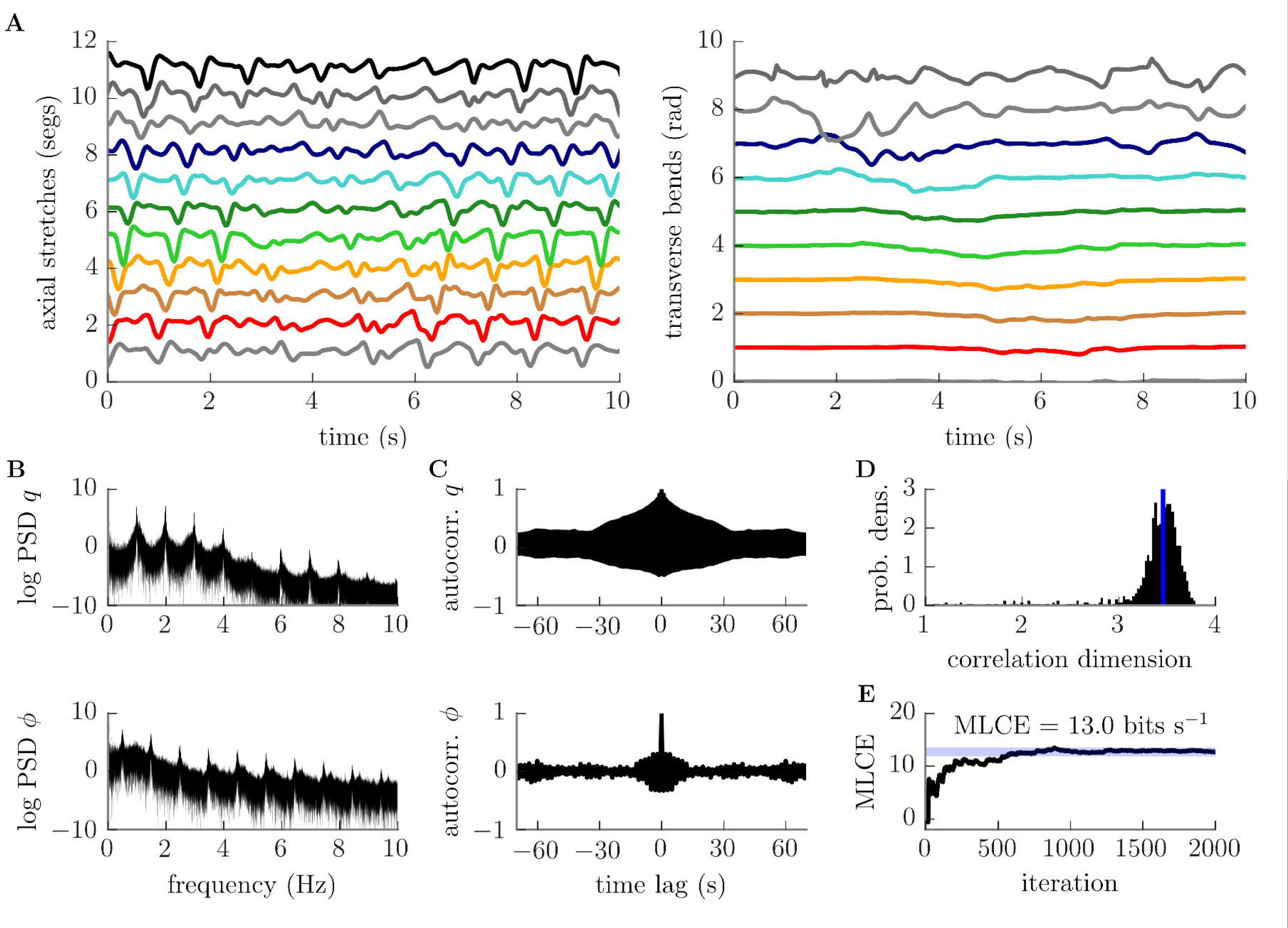
Dissipative planar motion is chaotic. A: representative segmental stretch (left) and bend (right) time series (see S3 Fig for corresponding neural state). Note the occurence of a large bend starting at ~ 1–2 seconds at the larva’s head, which appears to propagate backwards along the body while triggering a “transition” from forward to backward wave propagation at ~ 3.5 seconds. Forward wave propagation resumes at ~ 6 seconds. B: power spectra of the head stretch *q* (top) and bend *φ* (bottom) showing significant “noise” component. C: Autocorrelations of *q* and *$* rapidly decay. D: probability density of correlation dimension estimates for 1000 mechanical trajectories. The dimension of the system’s limit set is estimated as ~ 3.5 (median, vertical blue line). E: maximum Lyapunov characteristic exponent estimates converge to a positive value. All measures suggest the presence of deterministic chaotic dynamics. Parameters used to generate this figure are given in S5 Table.

The irregularity of the axial motion is reflected in the pattern of sensory neuron activation (S3 Fig). However, the mutual inhibitory interactions in our model act to filter this input, allowing only a small window of excitability within the central nervous system. As a result, interneuron and motor neuron activity appears fairly ordered, tracking and reinforcing axial compression waves.

We used four measures to assess whether our qualitative observation of irregular motion betrays the existence of deterministic chaos. First, we analysed the power spectrum of individual cooordinates (Fig 8B). The power spectra of all degrees of freedom showed consistent harmonic peaks along with a strong “noisy” non-harmonic component, a pattern consistent with chaos and incommensurate with (quasi)periodicity (Fig 8B shows data for head bending angle and stretch; similar data were obtained for other degrees of freedom, not shown). Next, we computed the autocorrelation of the same degrees of freedom. The autocorrelations of all degrees of freedom showed a periodic pattern with a peak at 0 seconds time lag followed by a rapid decay (Fig 8C). This is characteristic of oscillatory chaotic behaviour, and the exponential loss of information regarding initial conditions that chaotic systems display. We then estimated the correlation dimension (D_c_) of the limit set of our model’s dynamics. Note that we did not apply this measure to the conservative models in the previous section since the Liouville theorem rules out attracting limit sets for these systems. The distribution of correlation dimension estimates for our dissipative system across all 1000 trials is shown in Fig 8D. Estimates were clustered around ~ 3.5 (median dimension 3.46), with 93% of estimates lying in the range [3–4]. These results are suggestive of a limit set with fractal dimension, which is a signature of “strange” chaotic attractors. Finally, we computed an estimate of the maximal Lyapunov characteristic exponent (MLCE). As in the previous section, we estimated the MLCE for our system to be positive (~ 13bits s^−1^, Fig 8D), a very strong indicator of chaotic behaviour. All of these results point to the presence of a chaotic dynamical regime in our dissipative large amplitude model.

### Exploration emerges without decisions or stochasticity

As the coupled biomechanical and neuromuscular system produces both forward and backward peristalsis and lateral oscillations, each simulated larva will trace out a 2D trajectory over time. As shown in Fig 9A, the chaotic deformations characterised in the previous section caused the larvae to disperse across their two-dimensional substrate, following paths reminiscent of the real animal’s exploratory behaviour.

**Fig 9.**
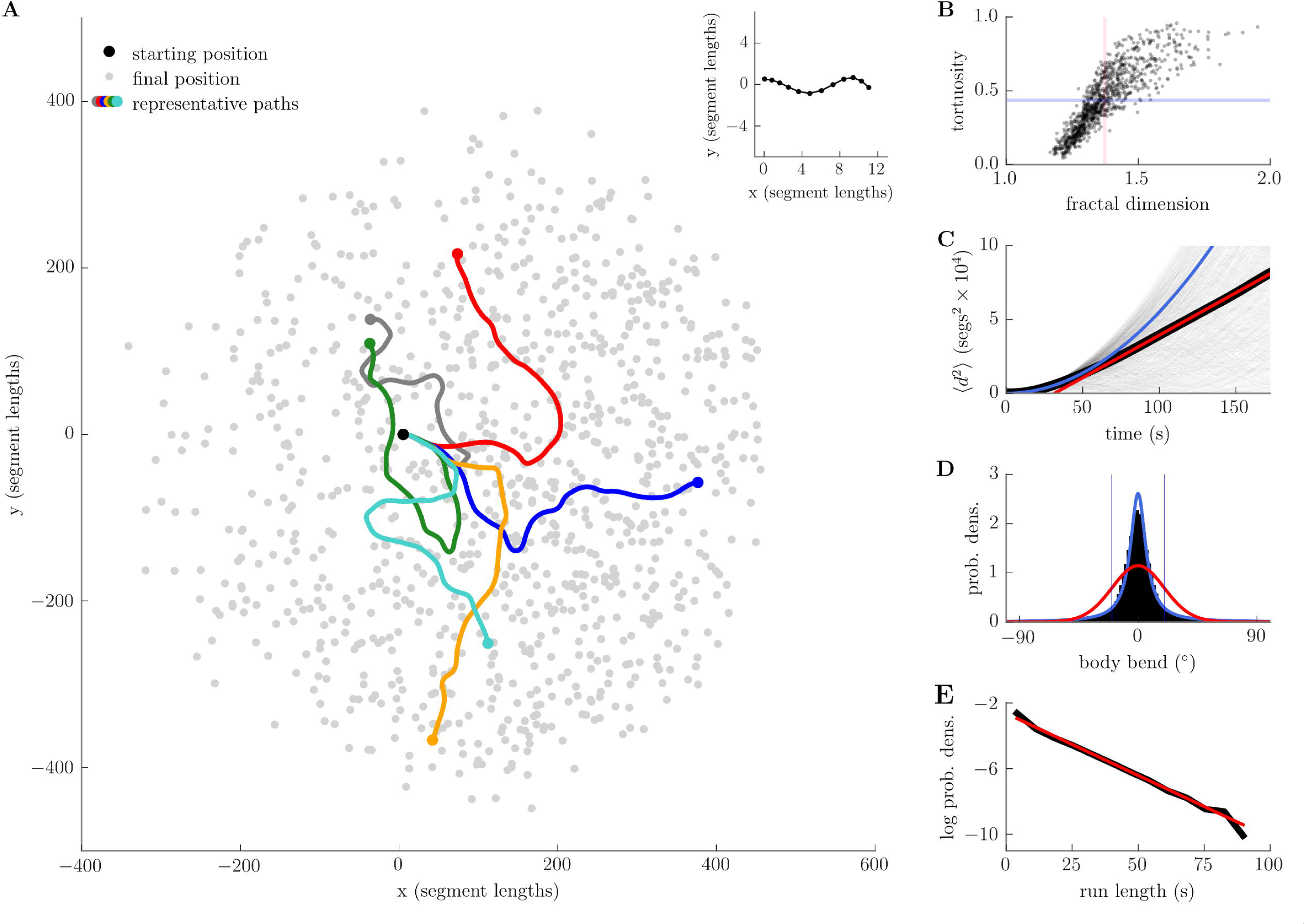
Deterministic exploration. A: dispersion of the centres of mass of 1000 simulated larvae, starting from almost identical mechanical initial conditions (overlayed in inset). B: tortuosity and fractal (box-counting) dimension for all 1000 paths indicate plane-filling behaviour (blue line = mean tortuosity, red line = mean dimension, see text; see S4 Fig for power law analysis of trajectory curvature and angular speed). C: mean-squared displacement (black line) shows transient quadratic growth (blue line) followed by asymptotic linear growth (red line, asymptotic diffusion constant ≈ 144segs^2^s^−1^; see also log-log plot, S5 Fig). D: distribution of body bends (black) with maximum likelihood von Mises (red) and wrapped Cauchy (blue) fits. E: distribution of run lengths with maximum likelihood exponential fit (red). Run lengths were calculated as duration between successive crossings of a threshold body bend (20°), indicated by blue lines in panel D. See S6 Fig for analysis of tail speed and head angular velocity. Parameters used to generate this figure are given in S5 Table.

To characterise the trajectories of our model, we first investigated them at a global level, based on the centre of mass (COM) trajectory of each simulated larva, computing the tortuosity and fractal dimension of the paths (Fig 9B) [54]. We defined our tortuosity measure as

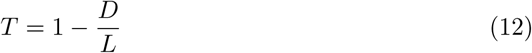

where *D* is the net displacement of the COM between initial and final times, and *L* is the total distance travelled by the COM along its path. Note that if the COM travels in a straight line between initial and final times we will have *D* = *L* so that *T* = 0. In the limit L → ∞ to, corresponding to the COM taking an extremely long path between its initial and final states, we have 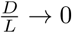so that T → 1. We calculated the mean tortuosity of our COM trajectories to be ⟨T⟩ = 0.43, with a variance of ⟨(T — ⟨T⟩⟩)^2^ = 0.05. The lowest (highest) tortuosity observed was *T* = 0.05 (*T* = 0.95).

We estimated the fractal dimension *D*_*b*_ of the COM trajectories using a box-counting algorithm. The minimum expected dimension *D*_*b*_ = 1 would correspond to curvilinear paths (e.g. straight line or circular paths), while the maximum expected dimension of *D*_*b*_ = 2 corresponds to plane-filling paths (e.g. brownian motion). We calculated the mean dimension of the COM trajectories to be ⟨*D*_*b*_⟩ = 1.37, with variance ⟨(*D*_*b*_ — ⟨*D*_*b*_⟩)^2^} = 0.01. The lowest (highest) path dimension observed was *D*_*b*_ = 1.17 (*D*_*b*_ = 1.95). We have plotted the tortuosity and fractal dimension of every path in Fig 9B. These results show that the trajectories of the model differed markedly from straight lines (tortuosity *T* > 0 and box-counting dimension *D*_*b*_ > 1), and displayed a good ability to cover the planar substrate (box-counting dimension 1< *D*_*b*_ < 2). We also note that our COM trajectories display the power-law relationship between angular speed and curvature reported by [55], with a scaling exponent (*β* ≈ 0.8) falling within the range reported for freely exploring larvae (S4 Fig).

We next investigated the rate at which the simulated larvae explored their environment. To do this, we calculated the mean-squared displacement (MSD) of the COM over time (Fig 9C). This is a standard measure used to characterise diffusion processes, and is defined as

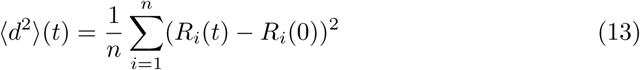

where *R*_*i*_(*t*) is the position of the *i*’th larva’s COM at time *t* and *n* = 1000 is the number of trials being averaged over. We observed an initial transient, lasting on the order of 10 seconds, during which the MSD grew as ~ *t*^2^ (blue line, S5 Fig), after which growth slowed and tended to ~ *t* (linear fit for *t* > 80 seconds shown by red line, Fig 9C and S5 Fig, *r*^2^ = 0.99, diffusion constant *D* = 144segs^2^s^−1^). The initial transient was not due to our particular initial conditions, since it remained even after discarding > 60s of initial data (not shown). These results therefore tell us that, although on long timescales our model appears to execute standard Fick diffusion or a Brownian random walk (linear growth of MSD), on short timescales the model’s behaviour is superdiffusive (approximately quadratic growth of MSD) [56,57]. This is in good agreement with observations of the real larva [6, 45]. The superdiffusive behaviour of the larva was previously explained in terms of a persistent random walk [6], in which the larva’s current and previous headings are highly correlated during straight runs so that the animal follows an approximately ballistic trajectory on short timescales. We believe that persistence effects arise in our model due to the finite time required for the deterministic chaotic dynamics to destroy information regarding initial conditions.

Larvae of different genetic backgrounds can show altered MSD growth rates relative to the wildtype animal [6,45]. We were able to increase or decrease the rate of exploration in our model by increasing or decreasing the transverse (bending) stiffness of the simulated larvae (not shown), a parameter which may in principle include both mechanical and neural components (see S5 Appendix).

We next calculated some other standard measures found in the larva literature. To do so, we built a two-segment representation of each simulated larva by drawing vectors from the tail extremity to the A2-A3 segment boundary (the tail vector, **T**), and from the A2-A3 boundary to the head extremity (the head vector, **H**). We then defined the body bend, 0, to be the angle between tail and head vectors, *θ* = atan(*H_y_/H_x_*) − atan(*T_y_/T_x_).* The head angular velocity *v* was computed as the cross-product of the head vector and the head extremity’s translational velocity 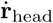 measured relative to the lab frame, 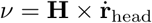, while the tail speed *v* was taken to be the magnitude of the tail extremity’s translational velocity 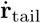 measured relative to the lab frame, 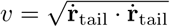. The tail speed and head angular velocity both show a strong oscillatory component, which can be seen in the time and frequency domains (S6 Fig). The power spectra of these variables contains considerable “noise” over a wide spread of frequencies, in accordance with the results of the previous section. The distribution of tail speeds for our model is bimodal, similar to that of the real larva [41]. The body bend angle was observed to be symmetrically distributed (Fig 9D), with roughly zero mean (⟨*θ*⟩ = 0.005), small variance (⟨(*θ* — ⟨*θ*⟩)^2^⟩ = 0.13), slight positive skew (*SK*(*θ*) = 0.23), and high excess kurtosis *KU*(*θ*) = 7.9. The kurtosis of our data precludes a good fit by the von Mises distribution (maximum likelihood estimate shown by red line in Fig 9D). Our data appears to be better fitted by a wrapped Cauchy distribution, though this overestimates the central tendency of our data (maximum likelihood estimate shown by blue line in Fig 9D). This analysis gives a quantitative measure of the rare large bending events mentioned at the beginning of the previous section (high excess kurtosis of the body bend distribution).

Finally, we computed a run-length distribution by setting a threshold body bend angle *θ*_*turn*_ = 20° (as in [13]) and calculating the length of time between successive crossings of this threshold. The distribution we obtained appears approximately linear on a log-linear plot (Fig 9E, linear fit *r*^2^ = 0.99 with slope λ = —0.075), and is better fit by an exponential than a power law distribution (maximum likelihood estimates, log likelihood ratio = 5281, *p* < 0.01). Together with our observation of asymptotic linear growth of MSD, the exponential distribution of run lengths suggests that the model can be considered to be effectively memoryless on long timescales [57]. This again agrees with the observed rapid loss of information from the system due to its chaotic dynamics, as quantified by the Lyapunov exponent and autocorrelation analysis of the previous section.

Ultimately, the analysis of our model supports a view of the larval exploratory routine as an (anomalous) diffusion process arising from the deterministic chaotic dynamics of the body. The model nervous system functions purely to recover these dynamics from the effects of frictive energy dissipation, and to ensure centre of mass motion, rather than explicitly directing exploration.

## Discussion

The intrinsic capabilities of an organism’s body determine the field of possibilities that neural circuits for behaviour can exploit. Here, by focusing first on the biomechanics of *Drosophila* larva, we find that its body already contains an inherent exploratory routine. This is demonstrated through a combined biomechanical and neuromuscular model that is the first to be able to generate both forward and backward peristalsis and turning, allowing 2D motion in the plane to be simulated. We show that, in the absence of friction, the body’s conservative mechanics alone supports both axial travelling waves and transverse standing waves. These are energetically coupled at larger amplitudes, such that no driving, sensing, or control of body bend is required for the system to start producing spontaneous coordinated bending motions. Frictional losses can be recovered, to maintain axial waves, by a neuromuscular system consisting of only simple local sensorimotor reflexes and long-range inhibitory interactions. This is sufficient to produce emergent crawling, reversal and turning that resembles larval exploratory behaviour, and which is chaotic in nature. At a population level, we observe a deterministic anomalous diffusion process in which an initial superdiffusive transient evolves towards asymptotic Fickian/Brownian diffusion, matching observations of real larvae [6,45]. We therefore propose that the role of biomechanical feedback in *Drosophila* larvae goes beyond the periphery of basic neuromuscular rhythms [39,40], to provide the essential “higher order” dynamics on which exploratory behaviour is grounded.

Most existing models of larval exploration abstract away from the mechanics underlying the production of runs and reorientations [4–6,8,11–13]. The larva is often described as executing a stochastic decision-making process which determines which state (running or turning) should be occupied, and when to initiate a change of behavioural state. In contrast, our model produces the entire exploratory routine without making any decisions (the transverse motion is neither sensed nor driven by the nervous system) nor introducing any stochastic process (neural or otherwise). Similarly, transient “switching” is seen to occur between forward and backward peristalsis even though there is no neural encoding or control of the direction of wave propagation. In other words, the body dynamics generate the basis of a chaotic exploratory routine which only needs to be amplified by the neural circuitry, making the search for underlying stochastic or state switching circuitry superfluous for this behaviour.

The work presented here also stands in contrast to previous models of larval peristalsis [38,40] and the prevailing hypotheses regarding this phenomenon [15,58] by eschewing any role for intrinsic neural dynamics. Such stereotyped and rhythmic locomotion is widely assumed to be the signature of a central pattern generator (CPG), that is, a neural circuit that intrinsically generates a rhythmic output, and thus determines a particular mechanical trajectory to be followed by the body [59–61]. However it is recognised that systems vary in the degree to which coordinated behaviour is independent of biomechanical and sensory feedback [61]. Indeed, evidence from studies employing genetic manipulations to disrupt sensory neuron input suggest that proprioceptive feedback is necessary for correct larval locomotive patterns [16,35–37,62]; although in some cases coordinated waves of forward and backward peristalsis can be produced, in both intact [16,35,62] and isolated VNC preparations [14,15], these are reported as abnormal with the most evident defects being time-dilation [15, 35, 62] and abnormal frequency in polarity changes [62].

In fact, our intent is not to adjudicate between the roles of intrinsically generated activation sequences vs. biomechanical feedback in this system, but rather to note that we should expect neural circuits of locomotion to adhere to the dynamical modes of the associated body, instead of working against them. Thus it should be unsurprising if these dynamics also exist (potentially in a latent form) in the neural circuitry. For example, a simple modification of the neural circuit presented here could produce instrinsic ‘peristaltic’ waves. Recall that the long-range global inhibition pattern in our model treats head and tail segments as ‘neighbouring’ nodes (see *Model specification and core assumptions - Neuromuscular system*). If local constant input or recurrent feedback were added to each segment, the circuit would then resemble a ring attractor [63–65] and a stable activity bump would be formed. Breaking the forward/backward symmetry of the circuit, e.g., by introducing asymmetric nearest-neighbour excitatory connections [66], would cause the activity bump to move along the network, giving rise to intrinsic travelling waves. This would complement any mechanical compression waves travelling through the body, i.e., remain consistent with the principles set out in this paper. Would such a network be a CPG? The answer is unclear. On the one hand, it would show spontaneous rhythmic activity in the absence of sensory input. On the other, sensory feedback would do much more than simply correct deviations from the CPG output or provide a “mission accomplished” signal [35]. Rather, feedback would play a crucial role in orchestrating motor output to ensure power flow into the body, consistently with its dynamical modes.

It is important to note that the emergence of rhythmic peristalsis and spontaneous turns in our model is not strongly dependent on the specific assumptions made in our mechanical abstraction. For example, the observation for small amplitude motions of sinusoidal axial travelling waves, along with transverse standing waves whose shapes match the experimentally observed “eigenmaggots” [48], is a direct result of the second-order Taylor series approximation of the model Hamiltonian (S2 Appendix). The small-amplitude model is thus non-unique, since many different mechanical models could have identical second-order approximations. Similarly, we expect that the deterministic chaotic behaviour derived from our conservative model for large amplitude motions will hold for other models of the larval body, given that it is conjectured that the majority of Hamiltonian systems are nonintegrable.

As a consequence of exploiting these mechanics, our model explains a wider range of behaviour than previous models, using a simpler nervous system. The properties included in the neuromuscular circuitry were derived from basic physical considerations,i.e., what was necessary and sufficient to produce exploration, rather than from known neuroanatomy or neurophysiology. However, it is useful to now examine what insights and predictions regarding this circuitry can be derived from our model.

Firstly, we consider the connections between segments. Unlike the model from [40], we did not require assymmetric connections to obtain forward (or backward) waves as these (and spontaneous switching between them) arise inherently in the mechanics. Rather, obtaining centre of mass motion of the entire body required the “ring attractor” layout of mutual inhibition between distant segments described above. The model thus predicts that motor output should be strongly inhibited (by signalling from other segments) the majority of the time, so that motor neurons only activate as the (mechanical) peristaltic wave passes through the corresponding body segment. This is in contrast to previous models which appealed only to local, nearest-neighbour inhibitory connections [38,40].

What might be the neural substrate for the proposed inhibition? There are two currently known intersegmental inhibitory pathways in the larva. GVLI premotor inhibitory neurons synapse onto motor neurons within the same segment but extend their dendritic fields several segments further anterior along the VNC. Accordingly, the GVLIs inhibit motor neurons at a late phase during the local motor cycle [22]. Our model predicts that there should be a larger set of GVLI-like neurons within each segment, with dendritic fields tiling distant segments. Although in our model the mutual inhibition is (for simplicity) arranged to act on all non-adjacent segments, we would in practice expect that active compression is actually spread across more segments [3,22] to transfer power to the body more efficiently (S3 Appendix), and this should be reflected in the inhibitory connection pattern. The second inhibitory pathway involves GDL inhibitory interneurons, which receive input from the excitatory premotor neuron A27h in the nearest posterior segment, and synapse onto A27h within the same segment while simultaneously disinhibiting premotor inhibitory neurons in distant segments [28]. Thus, GDL effectively produces both local and long-range inhibition of motor output. However, GDL receives axo-axonic connections from vdaA and vdaC mechanosensory cells within the same segment, so local inhibition is likely gated by sensory input. This would match our model, in which sensory activation within a segment should be sufficient to produce motor output when one of the neighbouring segments is active. We thus predict that simultaneous experimental suppression of GDL, GVLI, and all other long-range inhibition in the VNC should allow the propagation of several, concurrent locomotor waves in response to mechanical input.

Secondly, within a segment, our model highlights the importance of the timing of neuromuscular forces relative to body motion. Specifically, during locomotion, the larva’s muscles should act primarily as motors rather than as springs, brakes, or struts (see [33] for a discussion of these differences), and thus should activate in phase with the segmental stretch rate. This hypothesis could be tested by performing work-loop experiments, for which we predict the existence of a counterclockwise cycle in a plot of muscle force (potentially measurable by calcium imaging) over segment length during locomotion.

Can our model’s requirement that neurons sensing stretch-rate provide a direct excitatory connection to motor neurons, within the same segment, be mapped to identified pathways in the larva? One possible monosynaptic implementation of such a link are the dda mechanosensory cells which have been observed to make synapses onto aCC and RP2 motor neurons [23]. However, synapse counts show high variability both within and across individuals, so it seems unlikely to be a core component of the locomotor circuitry. A more promising candidate is the excitatory premotor interneuron A27h, which receives input from vpda and vdaC and sends bilaterally symmetric outputs to aCC and RP5 [28]. It is known that A27h activation is sufficient to activate downstream motor neurons, but it remains unknown whether proprioceptive sensory input is sufficient to activate A27h. Additionally, we hypothesise that AO2 (PMSI) interneurons [20], which have been recently shown to form an inhibitory sensory-motor feedback pathway between dbd mechanosensory cells and motor neurons [27], could play a role in filtering this signal to obtain the necessary stretch-rate activation independently of stretch. General models of mechanotransduction suggest that larval mechanosensory cells may be sensitive to both rate of stretch as well as absolute stretch, depending upon the mechanical properties of the sensory dendrites and the biophysics of the relevant mechanosensitive ion channels [67]. If PMSIs have a slow-activating, integrator dynamics that encodes stretch, while A27h activate quickly in response to proprioceptive sensory input to encode stretch and stretch-rate, the combined input to motor neurons would be only stretch-rate dependent excitation, as our model requires. This could explain the observation that optogenetic disturbance of PMSIs [20] slows the timescale of peristaltic waves, as the inclusion of absolute stretch in this feedback loop would produce muscle forces that not only counteract friction but also decrease the effective stiffness of the cuticle, slowing peristalsis (see S5 Appendix).

It is clear the real larval nervous system exhibits many complexities not reflected in our model, and likewise that the real larva performs many more behaviours than exploration. These include appropriate (directed) reactions to sensory stimuli such as stopping, withdrawal and reverse in response to touch stimuli [37]; and modulation of the frequency and direction of (large) turns in response to sensory gradients such as odour, heat or light [4,8–10,12,13,68–73] to produce positive or negative taxis. In a previous model of taxis [41] we have shown that by a continuous coupling of the amplitude of a regular lateral oscillation to the experienced change in stimulus strength in a gradient, a larva-like response to gradients can emerge, again without requiring active switching between states. In the current model, this could be effected by incorporating direct neuromuscular driving of bending degrees of freedom, since the real larva can likely use asymmetric activation of its lateralised muscles to produce active bending torques to influence the transverse motion. Alternatively, the degree of bend could be influenced indirectly by altering the stiffness and viscosity of segments, or their frictional interaction with the substrate. We note that the effective viscoelasticity of body segments can be neurally controlled by local reflex arcs (see S4 Appendix and [39]). Indeed, this could partially explain the experimental observation of increased bending on perturbation of a contralateral segmental reflex mediated by Eve+ interneurons [24]. The muscle activation caused by this reflex should produce bending torques which are proportional to current bend or bending rate, thus effectively modulating transverse stiffness or viscosity, respectively. Notably, in the taxis model of [41], it is not required that the descending signal that alters turn amplitude is lateralised, but rather that it has the right temporal coordination, which itself is naturally created by the interaction of body and environment.

The model presented in this paper does occasionally produce stops (cessation of peristalsis) during exploration, but this only occurs in concert with a large body bend (this stored transverse energy can subsequently and spontaneously restart the peristalsis); whereas in larva slowing, stopping and resumption of peristalsis (or transition from a stop to a large bend) can occur while the body is relatively straight [2,10]. As for ‘directed’ turning, this suggests that additional neural control might be needed to terminate or initiate movement in response to sensory stimuli. It is worth noting that our model predicts that peristalsis can be restarted by almost any small disturbance of the physical equilibrium provided the local feedback gain is high enough; similarly, lowering the gain means that energy losses due to friction are not compensated and the animal will stop. In general, we have found that altering assumptions about the sliding friction forces by which the model interacts with the substrate can often have unexpected and subtle effects on the motion produced, thus it would be interesting to further explore the functions provided by segmental lifting [3, 74], folding of the denticle bands (S1 Video), and extrusion of the mouth-hooks [3,75] during locomotion. In the more extreme case, larva are capable of burrowing through a soft substrate, and it is clear that a complex interaction of forces, mechanics, sensing and neural control must be involved that go well beyond the scope considered here.

## Supporting information

**S1. Fig. Coordinate system and substrate interaction schematics**. Internal coordinate system used to describe deformations of the larval body (left), and quantities used to describe substrate interaction (right). The friction force *F*_*friction*_ acting on the *i*’th segment boundary is directed opposite to that boundary’s velocity vector **v**_*i*_, and has a magnitude which depends only upon the direction *θ*_*i*_ of the velocity vector relative to a unit vector 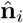 aligned with the local body axis (see text). Note that 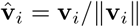 denotes a unit vector aligned with the boundary’s velocity vector.

**S2. Fig. Neural activation during peristalsis.** (from top to bottom) stretch rate, sensory neuron, interneuron, and motor neuron activation during forwards (left) and backwards (right) peristalsis.

**S3 Fig. Neural activation during planar locomotion.** (from top to bottom) stretch, stretch rate, sensory neuron, interneuron, and motor neuron activation during planar motion.

**S4 Fig. Relationship between path curvature and angular velocity.** Model data (grey points) compared to fit by a power law with scaling exponent *β* ≈ 0.8 (blue line, *r*^2^ ≈ 0.94).

**S5 Fig. log-log plot of mean-squared displacement.** Initial quadratic growth (blue line, slope=2) leads to asymptotic linear growth (red line, slope = 1).

**S6 Fig. tail speed v and head angular velocity *v* during planar motion. A:** representative time series for v and v. B: probability density of v and v across all 1000 trials. C: individual (faint) and mean (bold) power spectra of *v* and *v*

**S1 Video. Denticle bands fold into the larval cuticle during peristalsis.**

**S2 Video. Synthesis of travelling wave solutions from standing wave solutions.**

**S3 Video. Simulated larval exploratory behaviour.**

**S1 Appendix. Detailed model specification.**

**S2 Appendix. Detailed small-amplitude analysis.**

**S3 Appendix. A trade-off between power flow into the body and force on the centre of mass.**

**S4 Appendix. Modelling and analysis of head motion.**

**S5 Appendix. Effective body physics arising due to relationship of neuromuscular action to body motion.**

**S6 Appendix. Computer algebra and numerical methods.**

**S1 Table. Neural parameter values**. All segments are identical. Values given in larval units (seg = resting segment length, segmass = mass of a single segment boundary, nondim = dimensionless/nondimensional).

**S2 Table. Mechanical parameters for Fig 4 -** ***emergence of limit cycles for forward and backward locomotion in the dissipative, small-amplitude model***. All segments are identical. Values given in larval units (seg = resting segment length, segmass = mass of a single segment boundary, nondim = dimensionless/nondimensional).

**S3 Table. Mechanical parameters for Fig 6 -** ***changes in conservative head dynamics as amplitude of motion is increased***. Values given in larval units (seg = resting segment length, segmass = mass of a single segment boundary, nondim = dimensionless/nondimensional).

**S4 Table. Mechanical parameters for Fig 7 - *conservative planar motion of the body is chaotic at large amplitudes***. All segments are identical. Values given in larval units (seg = resting segment length, segmass = mass of a single segment boundary, nondim = dimensionless/nondimensional).

**S5 Table. Mechanical parameters for Fig 8** - ***dissipative planar motion is chaotic*** and Fig 9 - ***deterministic exploration***. Values given in larval units (seg = resting segment length, segmass = mass of a single segment boundary, nondim = dimensionless/nondimensional).

## Acknowledgments

We are grateful to Balazs Szigeti for providing us with experimental eigenmaggot data, and wish to thank Matthieu Louis and Richard Ribchester for their feedback on our manuscript.

